# Five-year (2017-2022) evolutionary dynamics of human coronavirus OC43 in southern France based on whole genome next-generation sequencing

**DOI:** 10.1101/2025.05.26.656003

**Authors:** Hikmat Houmadi, Céline Boschi, Lorlane Le Targa, PY Justine, Jean-Christophe Lagier, Lucile Lesage, Aurélie Morand, Bernard La Scola, Philippe Colson

**Affiliations:** Microbes Evolution Phylogeny and Infections (MEPHI), Aix-Marseille Univ. (AMU), 27 Bd Jean Moulin, 13005 Marseille, France; IHU Méditerranée Infection, 19-21 boulevard Jean Moulin, 13005 Marseille, France; Biosellal, 27 Chemin des Peupliers, 69570 Lyon, France; Assistance Publique-Hôpitaux de Marseille (AP-HM), 264 rue Saint-Pierre 13005 Marseille, France; Service de Pédiatrie générale, hôpital Timone, Assistance Publique-Hôpitaux de Marseille (AP-HM), 264 rue Saint-Pierre, 13005 Marseille, France; Service d’accueil des Urgences Pédiatriques, hôpital Nord, Assistance Publique-Hôpitaux de Marseille (AP-HM), chemin des Bourrely, 13015 Marseille, France

**Keywords:** Human coronavirus OC43, respiratory infections, genomics, mutations, lineages, recombination

## Abstract

HCoV-OC43 genomes and their evolution are scarcely studied worldwide and in France with only 361 genomes available as of October 2023. Here we implemented an in-house PCR amplification system to obtain retrospectively by next-generation sequencing then analyze HCoV-OC43 genomes for infections diagnosed with this virus in southern France between 02/2017 and 10/2022.

Multiplex PCR amplification using a set of in-house primers designed using the Gemi software was carried out on residues of HCoV-OC43 RNA-positive nasopharyngeal samples, before next-generation sequencing (NGS) using Illumina technology on a NovaSeq 6000 instrument. HCoV-OC43 genome assembly, bioinformatic analyses, and phylogeny reconstruction were then carried out using CLC Genomics, Mafft, BioEdit, Nextstrain, Nextclade, MEGA, iTOL and RDP4 softwares.

A total of 34 PCR primer pairs were designed for amplification then NGS of HCoV-OC43 genomes. A total of 185 genomes were obtained, 17, 79 and 89 belonging to genotypes G, J and K, respectively. These three genotypes circulated exclusively or co-circulated according to the year. A total of 303, 940, and 1,300 amino acid substitutions were detected in genotype G, J and K, respectively, compared with reference genomes of the same genotype dating back to 2017-2018. Possible recombinations were detected for 10 HCoV-OC43 genomes classified in genotypes K or J.

Overall, the present study more than doubled the set of HCoV-OC43 genomes available worldwide for the 2017-2022 period, and contributed to the monitoring of the HCoV-OC43 evolutionary dynamics.

## INTRODUCTION

Coronaviruses affect both humans and animals (Millet et al, 2020). Human coronaviruses are represented by endemic, or seasonal coronaviruses, including HCoV-OC43, HKU1, NL63 and 229E, as well as epidemic or pandemic coronaviruses including SARS-CoV, MERS-CoV and SARS-CoV-2 (Otieno et al, 2022; Martinez et al, 2021). Endemic coronaviruses are most often responsible for asymptomatic infections or mild upper respiratory diseases. HCoV-OC43 was in several studies the most frequently diagnosed human endemic coronavirus (Gaunt et al, 2010; Lau et al, 2006; Yip et al, 2016; Shah et al, 2022; Kong et al, 2021; Ye et al, 2023). It was reported to exhibit a seasonality of its incidence during winter and spring in western developed countries (Killerby et al, 2018; Rucinsky et al, 2020). In tropical countries, the seasonality of this virus is less clear (Otieno et al, 2022; Zhang et al, 2022; Al-Khannaq et al, 2016).

HCoV-OC43 was one the two first human coronaviruses that were discovered, in England during the 1960s, being first isolated from an organic tissue culture (McIntosh et al, 1967). Natural and intermediate hosts for HCoV-OC43 were described to be mice and cattle, respectively (Tang et al, 2022). This viral species was reported to share a common ancestor with a bovine coronavirus, and a bovine-to-human spillover was suspected to have occurred late during the 19^th^ century (Tang et al, 2022) while the most recent common ancestor of all genotypes was proposed to date back to the 1950s (Lau et al, 2011). HCoV-OC43 belongs to the genus *Betacoronavirus* (Tang et al, 2022). Its genome is approximately 30 kilobases in length, similar to that of the other coronavirus genomes. Two-thirds of this genome encode non-structural (Ns) proteins (Nsp1-16) and the remaining part encodes structural, and accessory proteins. The HCoV-OC43 genome notably encodes a haemagglutinin esterase, and accessory proteins include Ns2 and Ns12.9 (Zhang et al, 2015). HCoV-OC43 was reported to use 9-O-acetylated sialic acids as receptor to bind to host cells (Saunders et al, 2023; Hulswit et al, 2019).

Since the advent of next-generation sequencing (NGS), genomic data and studies particularly expanded for some viruses, but others remain neglected so far. Thus, while there are currently over 8 million SARS-CoV-2 genomes in GenBank (https://www.ncbi.nlm.nih.gov/genbank/) (Sayers et al, 2024) and 17 million in Gisaid (https://gisaid.org/) (Burki et al, 2023), only 361 (near) full-length HCoV-OC43 genomes (with a minimum coverage of 77%, i.e. a minimum size of 28,861 nucleotides) were available in GenBank as of October 2023. Moreover, only few studies were carried out on the genomic monitoring of this virus on large sample sets. Eleven genotypes (A-K) and an emerging lineage were described based on studies that analyzed genomes or the spike-encoding (S) gene (Zhang et al, 2022; Ye et al, 2023). Here we aimed to perform retrospectively the NGS of HCoV-OC43 genomes by implementing an in-house PCR amplification system, and to study the evolutionary dynamics of this virus for infections diagnosed between 2017 and 2022 in Southern France.

## MATERIALS AND METHODS

### Clincal samples

NGS of HCoV-OC43 genomes was performed retrospectively from remains of nasopharyngeal samples that had been diagnosed as HCoV-OC43 RNA-positive by real-time reverse-transcription (RT)-PCR (qPCR) using the FTD Respiratory pathogens 21 assay (FTD) (Fast Track Diagnosis, Luxembourg), after extraction by the KingFisher Flex system (Thermo Fisher Scientific, Waltham, MA, USA) according to the manufacter’s protocol, or by the FilmArray Respiratory panel 2 plus assay (BioFire) (Biomérieux, Marcy-l’Etoile, France). These clinical samples had been sent to our clinical microbiology laboratory at University and Public Hospitals of Marseille, Southeastern France, for the purpose of routine diagnosis of respiratory infections in the setting of clinical routine management, and they had been stored at −20°C or −80°C after their processing. A total of 1,007 nasopharyngeal samples were diagnosed HCoV-OC43-positive between 02/2017 and 10/2022 (69 months), and for 434 of them a sufficient volume was remaining on which the PCR amplification and NGS procedure could be applied.

### Primer design for PCR amplification of overlaping regions covering the whole viral genome

To design PCR primers, 220 complete HCoV-OC43 genomes were collected from the NCBI GenBank (https://www.ncbi.nlm.nih.gov/genbank/) and ViPR (https://www.bv-brc.org/view/Virus/10239) sequence databases. Then, these genomes were aligned using the MAFFT (Multiple Alignment using Fast Fourier Transform) software (Katoh et al, 2013), and the primers were designed using the GEMI software (Sobhy et al, 2012), an automated tool that enables quick and easy primer designs based on nucleotide conservation in a sequence alignment. GEMI was run by setting up amplicon size (500-2,600 nucleotides), amplicon overlap size (≥20 nucleotides), primer size (17-23 nucleotides), hybridization temperature (58-65°C), and also the maximal number of degenerated nucleotide positions (≤3). Primer specificity was then tested *in silico* using the NCBI BLAST tool (https://blast.ncbi.nlm.nih.gov/Blast.cgi) (Altschul et al, 1990). After primers were tested in simplex PCR assays and PCR were optimized using HCoV-OC43 RNA-positive cell culture supernatants, two sets of PCR primers were chosen, separated in two pools, to prevent the overlap of amplicons.

### PCR amplification of overlaping regions covering the whole genomes

Extracted RNA was amplified by standard RT-PCR using the SuperScript III One-Step RT-PCR Kit with Platinum Taq High Fidelity (Invitrogen, Life Technologies, Carlsbad, USA) according to the following protocol: 3 µL of RNA were added to 12.5 µL of 2X mix, 0.7 µL of polymerase, 1 µL of each primer pool, and 7.8 µL of pure water for a final volume of 25 µL. The PCR consisted of an initial RT step for 25 min at 50°C followed by an initial denaturation step for 2 min at 95°C, and by 40 PCR cycles including each 15 sec at 95°C, 45 sec at 58°C, and 150 sec at 70°C; a final elongation step consisted of 5 min at 70°C. Then, PCR amplicons were purified using the NucleoFast 96-well plate (Macherey Nagel, Hoerdt, France) for an elution volume of 40 µL. Eventually, PCR amplicons from the two PCR pools were mixed together as follows: 15 µL of pool no. 1 and 25 µL of pool no. 2.

### Next-generation sequencing

To test the PCR primers for the multiplex amplification system separately and pooled on HCoV-OC43 RNA-positive cell culture supernatants, we performed NGS using the Oxford Nanopore Technology with the Ligation Sequencing Kit (SQK-LSK109) then library sequencing on a GridION instrument after deposit on a SpotON flow cell Mk I, R9.4.1, following to the manufacturer’s instructions (Oxford Nanopore Technologies, Oxford, UK). Thereafter, when the ‘Artic-like’ system was deemed optimal, we performed NGS on RNA extracts obtained from remains available for nasopharyngeal samples that had been diagnosed as HCoV-OC43 RNA-positive in our laboratory, using the Illumina technology on a NovaSeq 6000 instrument. At this stage, NGS used the COVIDSeq protocol (Illumina Inc., San Diego, CA, USA), but by replacing the COVID-19 ARTIC PCR primers by PCR primers designed here, and using the conditions we previously set up. A classic loading procedure on a SP flow cell was carried out according to the NovaSeq-XP workflow and a previously described procedure (Papa Mze et al, 2023), with a reading of 2 x 50 nucleotides.

### Bioinformatic analyses

A workflow was developed using the CLC genomics workbench software v.7, consisting of a quality control step that included a trimming with a minimum of 15 nucleotides in the reads, followed by consensus genome generation by a mapping step of NGS reads against a reference genome using as parameters a coverage ≥80% and a nucleotide identity ≥90%. For the phylogenetic analysis, consensus genome sequences were aligned using the Mafft software (Katoh et al, 2013) with default parameters, then the tree was built using Iqtree2 (Minh et al, 2020)with the maximum likelihood method and 1,000 boostrap replicates, and it was visualized and annotated using the iTOL software (Letunic et al, 2007). To search for amino acid and nucleotide mutations, the Nextclade and Nextstrain tools (https://nextstrain.org) (Aksamentov et al, 2021)were modified to be used for HCoV-OC43 sequences, and analyses were conducted separately according to the genotype. Reference genomes GenBank accession no. MN026164.1 (dating back to 2018) for genotype G, OK318939.1 (dating back to 2018) for genotype J, and MW532118 (dating back to 2017) for genotype K were used. Viral genes were classified into “structural”, “informational”, “other non-structural” and “accessory” categories as in a previous study (Colson et al, 2023); the hemagglutinin esterase (HE) encoding gene and the N2 part of the nucleocapsid encoding gene were added in the structural gene category, while the category of accessory genes included Ns2 and Ns12.9. Finally, possible recombination events into HCoV-OCV43 genomes were searched out using the RDP4.101 software (Martin et al, 2015)on genomes with at least 95% coverage of full-length reference genomes. This software concurrently performs seven tests including RDP, Geneconv, Bootscan, Maxchi, Chimaera, SiSscan and 3Seq. A potential recombination was defined by ≥5 positive tests with a p value ≤0.001, as previously reported to study recombination events in the case of the West Nile virus (Mencattelli et al, 2023).

## RESULTS

### Primer design for PCR amplification of overlaping regions covering the whole genomes

Of a total of 220 HCoV-OC43 genomes collected from GenBank, 165 were selected for PCR primer design. A total of 200 PCR primer pairs were obtained by GEMI and 34 were eventually retained. These PCR primers had a maximum of two degenerated nucleotides, their size varied between 17 and 23 nucleotides and the expected sizes of the amplicons varied between 500 and 2,600 base nucleotides. Melting temperatures varied between 58°C and 65°C and the PCR amplicons overlapped by at least 20 nucleotides. These 34 primers were tested individually and their efficacy was verified by visualizing the presence of bands on a 1.5% agarose gel after DNA staining and electrophoretic migration. PCR optimization was carried out by increasing the primer concentration in case of weak bands on the agarose gel. Eventually, the 34 PCR primers were mixed into two different pools of 17 primer pairs each to avoid overlaps between the PCR amplicons, with primer concentrations ranging between 10 and 15.3 µM (Table 1).

**Table 1.**
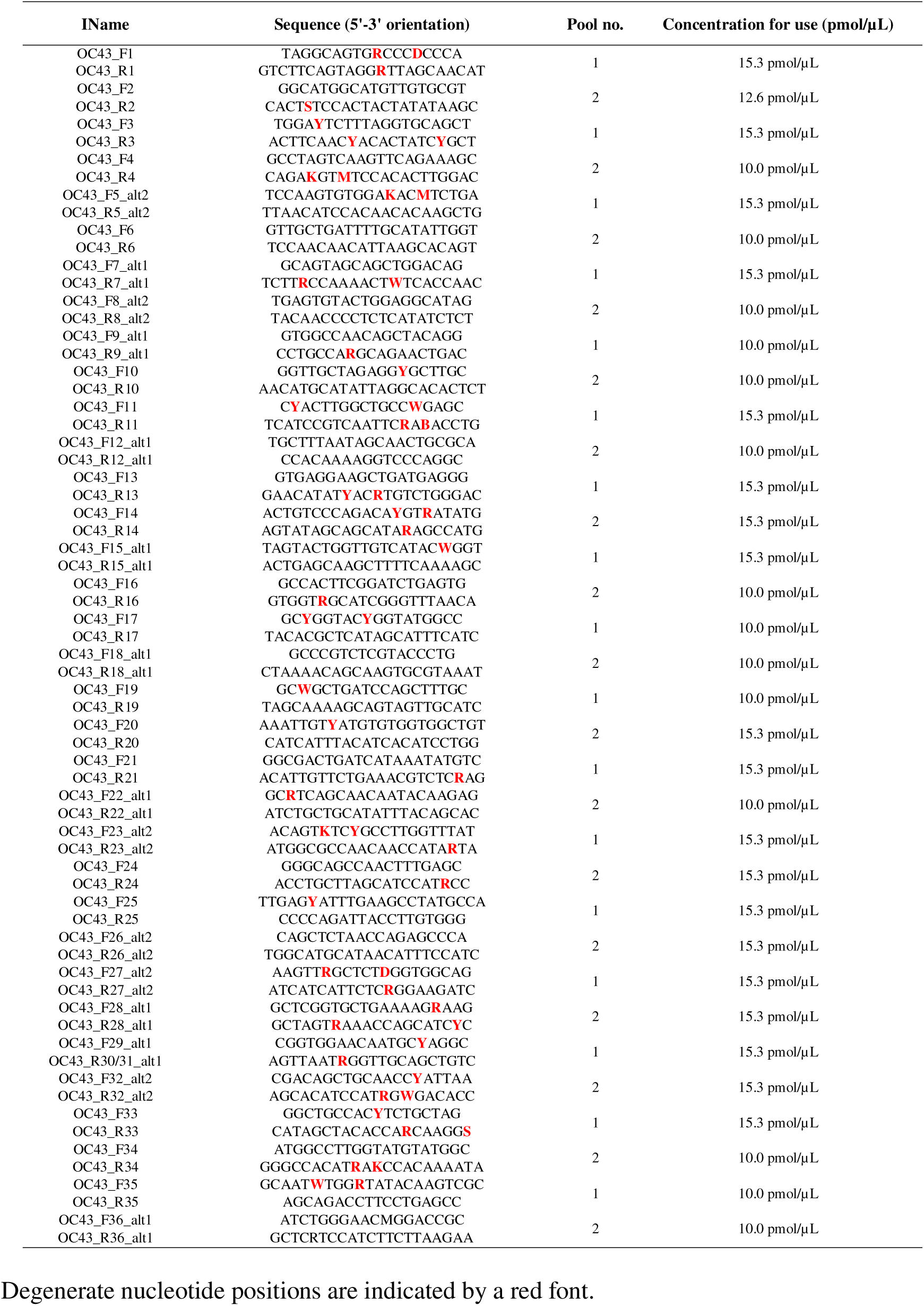
Sequences of primers used in the multiplex PCR amplification system and conditions of use.

### Next-generation sequencing

After amplification of HCoV-OC43 genomes using our in house multiplex PCR system, NGS was performed on 434 HCoV-OC43 RNA-positive nasopharyngeal samples with the Illumina technology on a NovaSeq 6000 instrument. A total of 185 HCoV-OC43 genomes were recovered according to our chosen parameters including a minimum of 80% coverage of full-length reference genomes and a minimum of 3X as NGS depth relatively to the references. The 185 nasopharyngeal samples from which they were obtained had been collected between 06/02/2017 and 16/07/2022 (over a period of 66 months). The qPCR cycle threshold values (Ct) were available for 96 samples and ranged between 11 and 25 with a mean (± standard deviation) of 17.8 ± 3.3 Ct. The total number of NGS reads per sample that were mapped onto the reference genome ranged between 5,675 and 9,917,033, with a mean of 3,569,789±2,943,862 reads, and mean coverage was of 94.7±5.2%.

### Phylogeny reconstruction and temporal distribution of genotypes

Phylogeny reconstruction included HCoV-OC43 genome sequences obtained here as well as genomes of the different genotypes from datasets obtained in previous studies (Ye et al, 2023). Only boostrap values ≥70% were displayed on the tree. Viral genome recovered here were clustered into three genotypes, namely G, J and K, with 17, 79 and 89 sequences, respectively (Figure 1). Mean nucleotide diversity between sequences obtained here was 91.9±6.3 for genotype G, 93.2±5.1 for genotype J, and 91.8±5.3 for genotype K. Regarding the temporal distribution of the incidence of the different genotypes (Figure 2), a single genotype circulated during some periods of time, such as for the case of genotype K between November 2020 and October 2021 (Figure 2). In constrast, during other periods, several genotypes co-circulated, as for the case of genotypes J and K in 2020, or of all three genotypes G, J and K in 2018 and in 2022.

**Figure 1.**
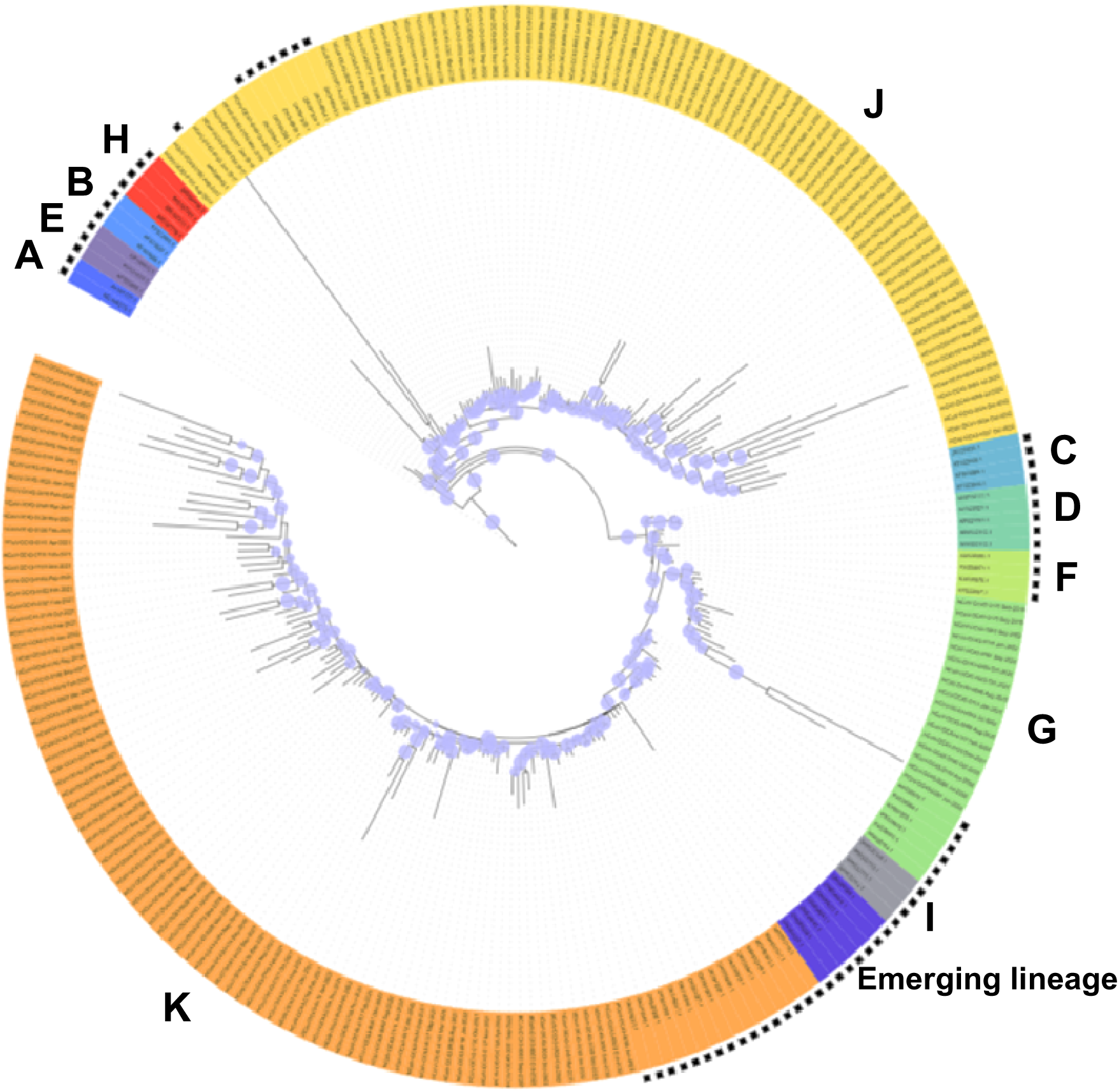
Phylogenetic tree based on HCoV-OC43 genomes obtained in the present study and from GenBank Boostrap values ≥70% were indicated at nodes on the tree by a light blue circle. Genomes recovered from GenBank (https://www.ncbi.nlm.nih.gov/genbank/) (Sayers et al, 2024) and not obtained here are indicated by a black asterisk.

**Figure 2.**
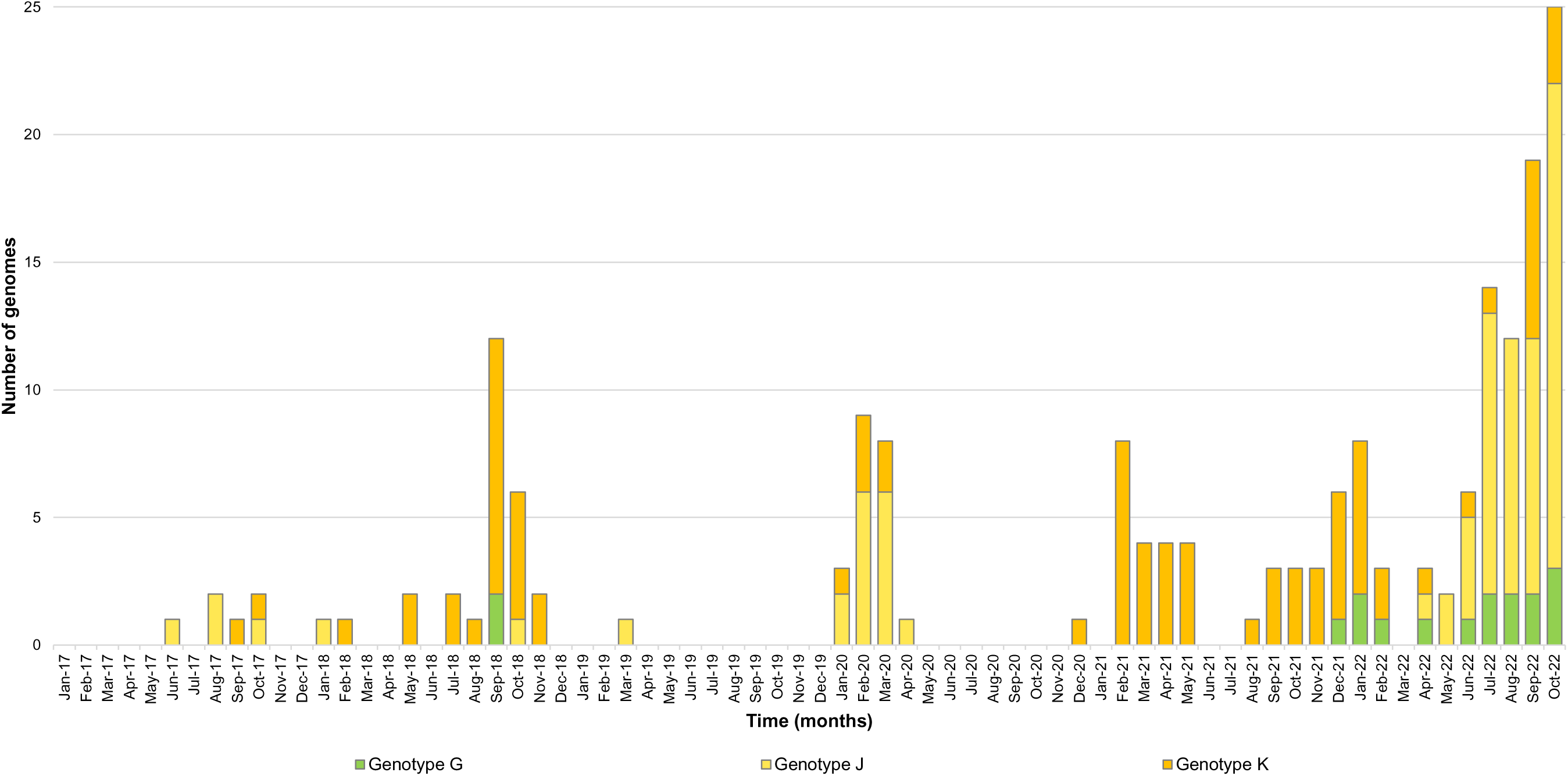
Temporal distribution of genotypes for HCoV-OC43 genomes obtained in the present study

### Mutation patterns

The patterns of mutations within genomes were analyzed for each viral genotype separately. Regarding genotype G genomes, 991 nucleotide substitutions were detected compared with the reference genome GenBank accession no. MN026164.1 dating back to 2018. To gain a better picture of the distribution of these mutations, we classified the genes into “structural”, “informative”, “other non-structural” and “accessory” categories. In the informational gene category, Nsp13 that encodes the helicase was the most affected gene, with 46 substitutions per 1,000 nucleotides and Nsp15 that enccodes an endoribonuclease was the least affected, with 5 substitutions per 1,000 nucleotides (Table 2a). In the structural gene category, the envelope gene showed the greatest diversity with 106 substitutions per 1,000 nucleotides and the matrix encoding gene was the least affected with 4 substitutions per 1,000 nucleotides. A total of 303 amino acid substitutions were detected relatively to the reference genome MN026164.1 (Table 2a; Figure 3a). In the informational gene category, Nsp13 was the most affected with 44 amino acid substitutions per 1,000 nucleotides and Nsp8 exhibited no amino acid substitution. In the structural gene category, the most affected gene was the envelope gene and the least affected was the matrix gene. We observed that 41 different amino acid substitutions were encoded by ≥10 of the genotype G genomes. The genes harboring these mutations were Nsp2, Nsp3, Nsp4, Nsp12, Nsp13, Nsp14, and Nsp16 regarding non-structural genes, and HE, S, N and N2 regarding structural genes. A majority of these mutations were located in the spike gene and were present in the majority of the genomes analyzed here. One mutation (S507G) was located in the receptor binding domain (RBD) of the spike protein that was predicted to span amino acids 339-549 (Lau et al, 2011). The other genes including Nsp1, Nsp5, Nsp6, Nsp7, Nsp8, Nsp9, Nsp10, Nsp11, Nsp15, M, E and the accessory genes were considerably conserved.

**Figure 3.**
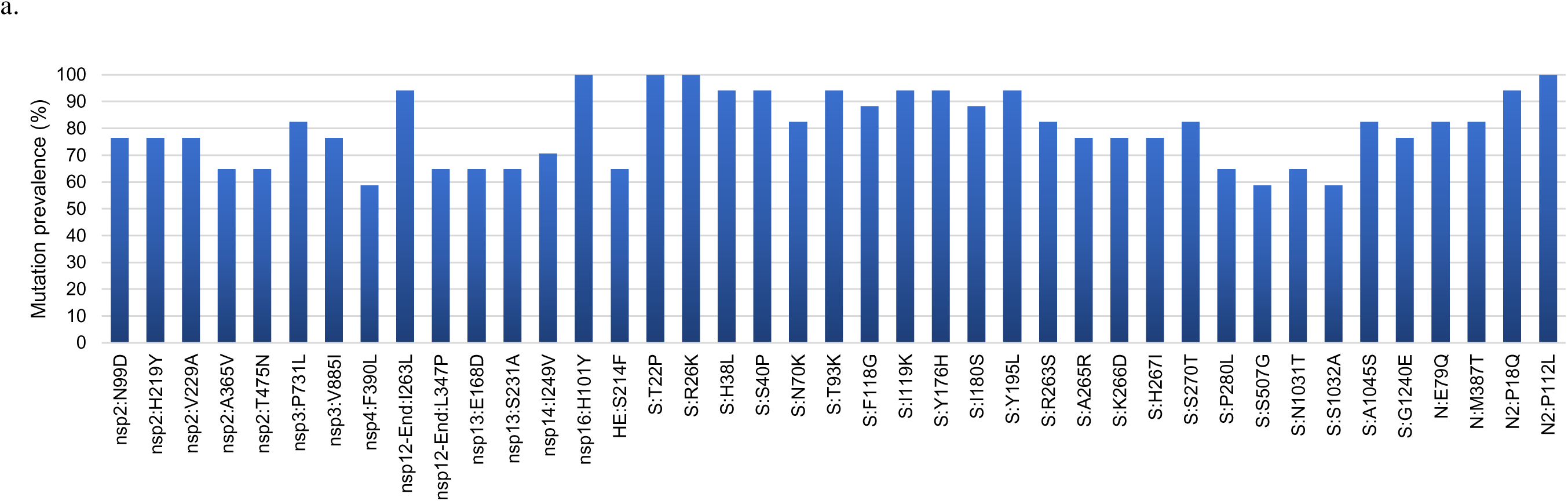
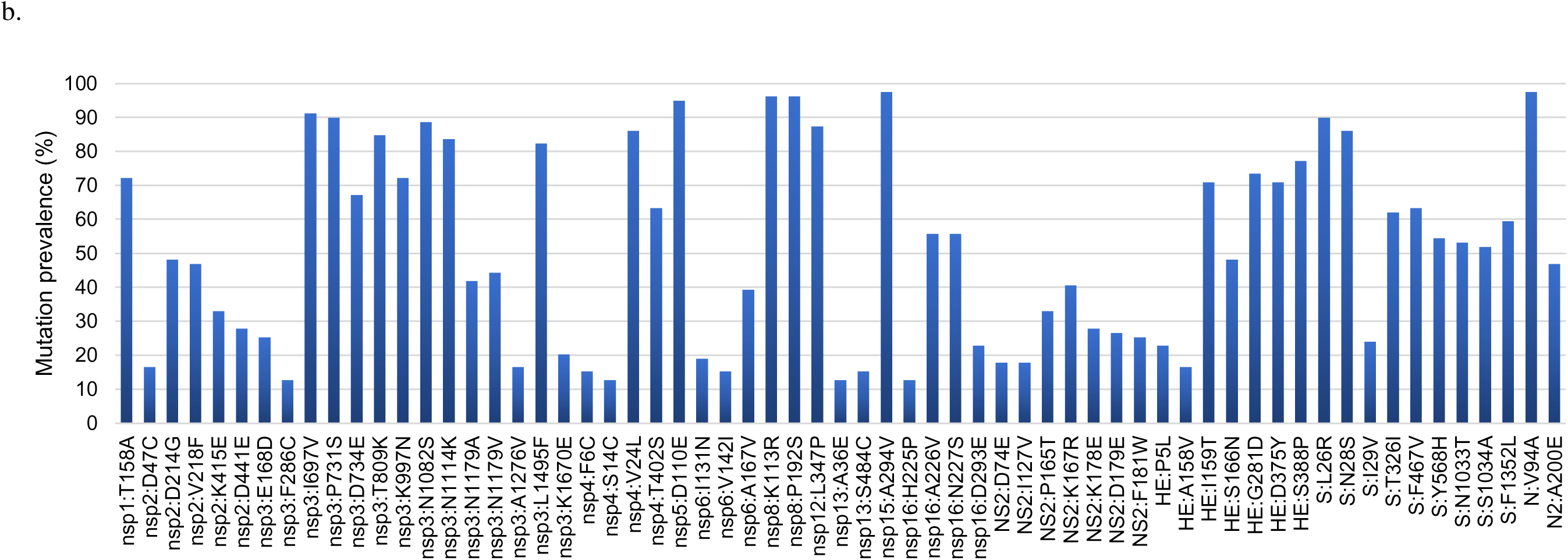
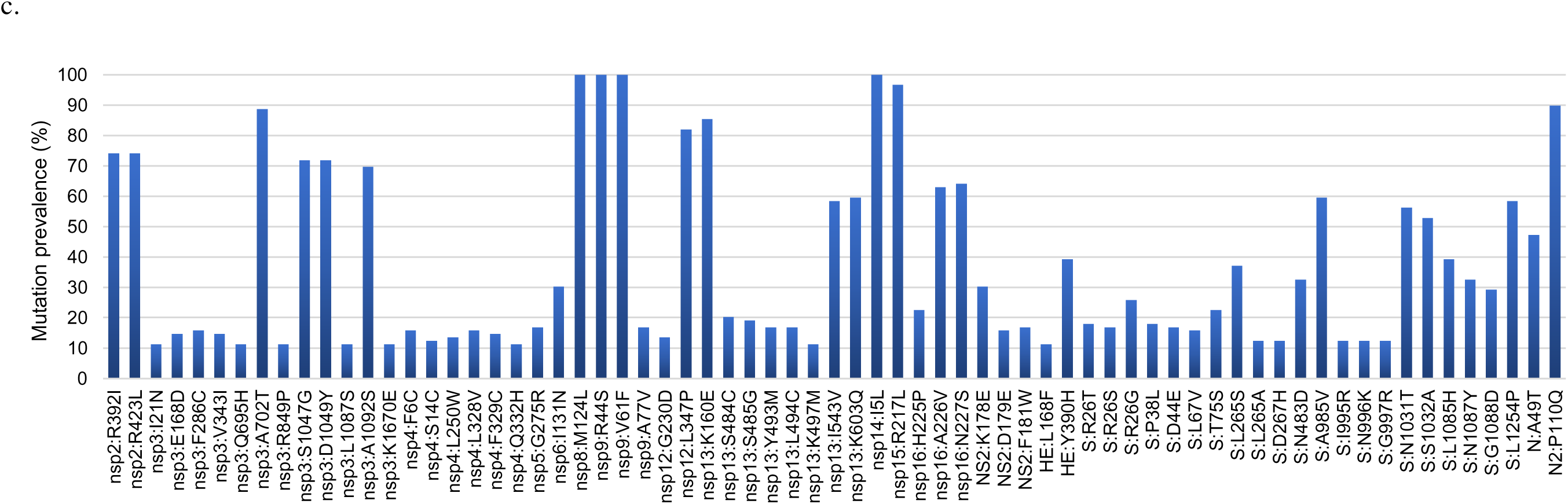
Distribution along HCoV-OC43 genomes of the three genotypes G (a), J (b), and K (c) obtained here of the prevalence of amino acid substitutions relatively to reference genomes References genomes were genomes GenBank (https://www.ncbi.nlm.nih.gov/genbank/) (Sayers et al, 2024) Accession no. MN026164.1, OK318939.1 and MW532118.1 for genotype G, J and K, respectively. HE, hemagglutinine esterase; N, nucleocapsid; NS, non-structural; nsp, non-structural protein; S, spike.

**Table 2.** Frequency of nucleotide and protein mutations in HCoV-OC43 genotype G (a), J (b) and K (c)

| Gene | Size<br>(nucleotides) | Number of<br>mutations per<br>gene | Number of<br>mutations per<br>1,000 nucleotides | Size<br>(amino acids) | Number of<br>mutations per<br>protein | Number of mutations<br>per 1,000 amino acids |
| --- | --- | --- | --- | --- | --- | --- |
| Nsp1 | 738 | 1 | 1 | 222 | 1 | 4 |
| Nsp2 | 1,815 | 71 | 39 | 587 | 15 | 25 |
| Nsp3 | 5,697 | 130 | 23 | 2,029 | 36 | 17 |
| Nsp4 | 1,488 | 107 | 72 | 496 | 35 | 70 |
| Nsp5 | 909 | 8 | 9 | 303 | 2 | 6 |
| Nsp6 | 861 | 4 | 4 | 287 | 4 | 14 |
| Nsp7 | 267 | 2 | 7 | 92 | 1 | 11 |
| Nsp8 | 591 | 4 | 6 | 194 | 0 | 0 |
| Nsp9 | 330 | 3 | 9 | 110 | 1 | 9 |
| Nsp10 | 411 | 5 | 12 | 137 | 1 | 7 |
| Nsp11 | 42 | 0 | 0 | 14 | 0 | 0 |
| Nsp12 | 2,783 | 82 | 29 | 928 | 35 | 37 |
| Nsp13 | 1,809 | 85 | 46 | 603 | 27 | 44 |
| Nsp14 | 1,563 | 13 | 8 | 521 | 2 | 4 |
| Nsp15 | 1,125 | 6 | 5 | 374 | 4 | 10 |
| Nsp16 | 897 | 22 | 24 | 299 | 12 | 40 |
| NS2 | 837 | 65 | 77 | 278 | 25 | 90 |
| HE | 1,284 | 58 | 45 | 427 | 17 | 40 |
| S | 4,077 | 176 | 43 | 1,358 | 57 | 42 |
| NS12.9 | 330 | 6 | 18 | 109 | 2 | 18 |
| E | 255 | 27 | 106 | 84 | 12 | 143 |
| M | 693 | 3 | 4 | 230 | 1 | 4 |
| N | 1,347 | 36 | 26 | 448 | 8 | 18 |
| N2 | 348 | 5 | 14 | 115 | 5 | 43 |

**b**
| Gene | Size<br>(nucleotides) | Number of<br>mutations per<br>gene | Number of<br>mutations per<br>1,000 nucleotides | Size<br>(amino acids) | Number of<br>mutations per<br>protein | Number of mutations<br>per 1,000 amino acids |
| --- | --- | --- | --- | --- | --- | --- |
| Nsp1 | 738 | 101 | 137 | 222 | 28 | 126 |
| Nsp2 | 1,815 | 157 | 86 | 587 | 58 | 99 |
| Nsp3 | 5,697 | 329 | 57 | 2,029 | 107 | 52 |
| Nsp4 | 1,488 | 168 | 113 | 496 | 58 | 117 |
| Nsp5 | 909 | 31 | 34 | 303 | 9 | 29 |
| Nsp6 | 861 | 70 | 81 | 287 | 25 | 87 |
| Nsp7 | 267 | 10 | 37 | 92 | 4 | 43 |
| Nsp8 | 591 | 11 | 18 | 194 | 4 | 20 |
| Nsp9 | 330 | 5 | 15 | 110 | 4 | 36 |
| Nsp10 | 411 | 4 | 9 | 137 | 0 | 0 |
| Nsp11 | 42 | 0 | 0 | 14 | 0 | 0 |
| Nsp12 | 2,783 | 244 | 87 | 928 | 47 | 50 |
| Nsp13 | 1,809 | 210 | 116 | 603 | 54 | 89 |
| Nsp14 | 1,563 | 45 | 28 | 521 | 8 | 15 |
| Nsp15 | 1,125 | 61 | 54 | 374 | 18 | 48 |
| Nsp16 | 897 | 57 | 63 | 299 | 20 | 67 |
| NS2 | 837 | 289 | 345 | 278 | 115 | 413 |
| HE | 1,272 | 313 | 246 | 423 | 127 | 300 |
| S | 4,083 | 713 | 174 | 1,360 | 189 | 139 |
| NS12.9 | 330 | 11 | 33 | 109 | 3 | 27 |
| E | 267 | 12 | 45 | 88 | 7 | 79 |
| M | 693 | 48 | 69 | 230 | 18 | 78 |
| N | 1,344 | 74 | 55 | 447 | 22 | 49 |
| N2 | 624 | 36 | 57 | 207 | 15 | 72 |

c.
| Gene | Size<br>(nucleotides) | Number of<br>mutations per<br>gene | Number of<br>mutations per<br>1,000 nucleotides | Size<br>(amino acids) | Number of<br>mutations per<br>protein | Number of mutations<br>per 1,000 amino acids |
| --- | --- | --- | --- | --- | --- | --- |
| Nsp1 | 738 | 64 | 86 | 222 | 14 | 63 |
| Nsp2 | 1,815 | 246 | 135 | 587 | 82 | 139 |
| Nsp3 | 5,697 | 672 | 118 | 2,029 | 182 | 89 |
| Nsp4 | 1,488 | 313 | 210 | 496 | 106 | 213 |
| Nsp5 | 909 | 105 | 115 | 303 | 34 | 112 |
| Nsp6 | 861 | 85 | 98 | 287 | 25 | 87 |
| Nsp7 | 267 | 11 | 41 | 92 | 6 | 65 |
| Nsp8 | 591 | 46 | 78 | 194 | 22 | 113 |
| Nsp9 | 330 | 21 | 63 | 110 | 10 | 91 |
| Nsp10 | 411 | 13 | 31 | 137 | 5 | 36 |
| Nsp11 | 42 | 0 | 0 | 14 | 0 | 0 |
| Nsp12 | 2,783 | 315 | 113 | 928 | 58 | 62 |
| Nsp13 | 1,809 | 300 | 166 | 603 | 78 | 129 |
| Nsp14 | 1,563 | 73 | 46 | 521 | 32 | 61 |
| Nsp15 | 1,125 | 78 | 69 | 374 | 25 | 67 |
| Nsp16 | 897 | 103 | 115 | 299 | 39 | 130 |
| NS2 | 837 | 381 | 455 | 278 | 138 | 496 |
| HE | 1,284 | 318 | 247 | 427 | 114 | 267 |
| S | 4,077 | 833 | 204 | 1,358 | 228 | 168 |
| NS12.9 | 330 | 31 | 94 | 109 | 11 | 101 |
| E | 255 | 49 | 192 | 84 | 15 | 178 |
| M | 693 | 78 | 112 | 230 | 20 | 87 |
| N | 1,347 | 117 | 87 | 448 | 38 | 85 |
| N2 | 624 | 57 | 91 | 207 | 18 | 87 |

HE, hemagglutinine esterase; N, nucleocapsid; NS, non-structural; nsp, non-structural protein;
S, spike.

Regarding genotype J genomes, their mutation patterns were analyzed relatively to reference genome OK318939.1 that dates back to 2018. In the informational gene category, the gene with the greatest nucleotide diversity was Nsp13, and the one with the lowest diversity was Nsp10, with 116 and 9 substitutions per 1,000 nucleotides, respectively (Table 2b). In the structural gene category, the gene most affected was the one encoding HE with 246 substitutions per 1,000 nucleotides, followed by the spike gene with 174 substitutions per 1,000 nucleotides, and the gene the least affected was the envelope gene with 45 substitutions per 1,000 nucleotides. A total of 940 amino acid substitutions were observed to be encoded by the genotype J genomes obtained here relatively to genome OK318939.1 (Table 2b; Figure 3b). In the informational gene category, Nsp13 was the most mutated with 89 amino acid substitutions per 1,000 nucleotides and Nsp10 was the least mutated with no amino acid substitution. In the informational gene category, Nsp13 was the most affected with 89 amino acid substitutions per 1,000 nucleotides and Nsp10 was the least affected with no amino acid substitution. In the structural gene category, the HE encoding gene was the one exhibiting the greatest amino acid diversity, while the spike and nucleocapsid genes were those the less mutated. In the accessory gene category, Ns2 showed the greatest amino acid diversity with 413 amino acid substitutions per 1,000 nucleotides. Overall, 64 amino acid substitutions were encoded by ≥10 genotype J genomes. A great number of mutations were observed in non-structural genes, the most mutated of which was Nsp3. Structural gene products with amino acid changes were HE, spike and nucleocapsid. One mutation (F467V) was located in the spike RBD. There were also mutations in the Ns2 accessory gene. Finally, the most conserved gene products were Nsp7, Nsp10, Nsp11, Nsp14, M, E and the other accessory proteins apart from Ns2 (Figure 3b).

Regarding genotype K genomes, mutations were analysed relatively to genotype K reference genome MW532118.1 that dates back to 2017. The gene with the greatest nucleotide diversity in the informational gene category was Nsp13 and the gene with the lowest diversity was Nsp10, with 166 and 31 substitutions per 1,000 nucleotides, respectively. In the structural gene category, the HE gene was the one with the greatest diversity with 247 substitutions per 1,000 nucleotides, followed by the spike gene with 204 substitutions per 1,000 nucleotides, while the nucleocapsid gene was the one with the lowest diversity with 87 substitutions per 1,000 nucleotides (Table 2c). Ns2 was the accessory gene with the greatest diversity with 455 substitutions per 1,000 nucleotides. Besides, a total of 1,300 amino acid substitutions were detected in the genomes obtained in the present study relatively to reference genome MW532118.1. In the informational gene category, the greatest amino acid diversity was observed for Nsp16 and Nsp13, with 130 and 129 amino acid substitutions per 1,000 nucleotides, respectively. Nsp10 was the gene the less affected by mutations in this category. In the structural gene category, the HE gene was the gene with the greatest diversity and the nucleocapsid gene was the gene with the lowest diversity, with 267 and 85 amino acid substitutions per 1,000 nucleotides, respectively. Ns2 was the accessory gene with the greatest diversity (Table 2c; Figure 3c). Overall, 68 amino acid changes were found in ≥10 genomes. They were distributed in almost every gene, in both non-structural and structural genes, including in Nsp2, Nsp3, Nsp4, Nsp5, Nsp6, Nsp8, Nsp9, Nsp12, Nsp13, Nsp14, Nsp15, Nsp16, HE, S, N, N2 and the accessory gene Ns2. One mutation (N483D) was located in the spike RBD. The most conserved proteins were Nsp1, Nsp7, Nsp10, Nsp11, M, E and the other accessory proteins apart from Ns2.

### Genome recombination events

Recombination events were searched for genomes with ≥95% coverage using the RDP4 software. Potential recombination events were detected for genomes retrieved from 10 samples. Of these 10 genomes, 6 belonged to genotype J and 4 to genotype K (Table 3). The genotypes of major-minor parental sequences were K-J and J-K in three cases each; and K-I, K-E, C-F and I-J in one case each. Therefore, these putative recombinant genomes involved parental genomes of genotypes not found here as non-recombinant genomes, including C, E, F, and I. The breakpoints differed depending on the genome, being located in different genes. They involved the 5’untranslated region (UTR) in two cases; the 3’UTR in two cases; the Ns2, S, and HE genes in three cases each, the Nsp12, Nsp13 and Nsp16 genes in two cases each; and the N gene in one case (Table 3).

**Table 3.**
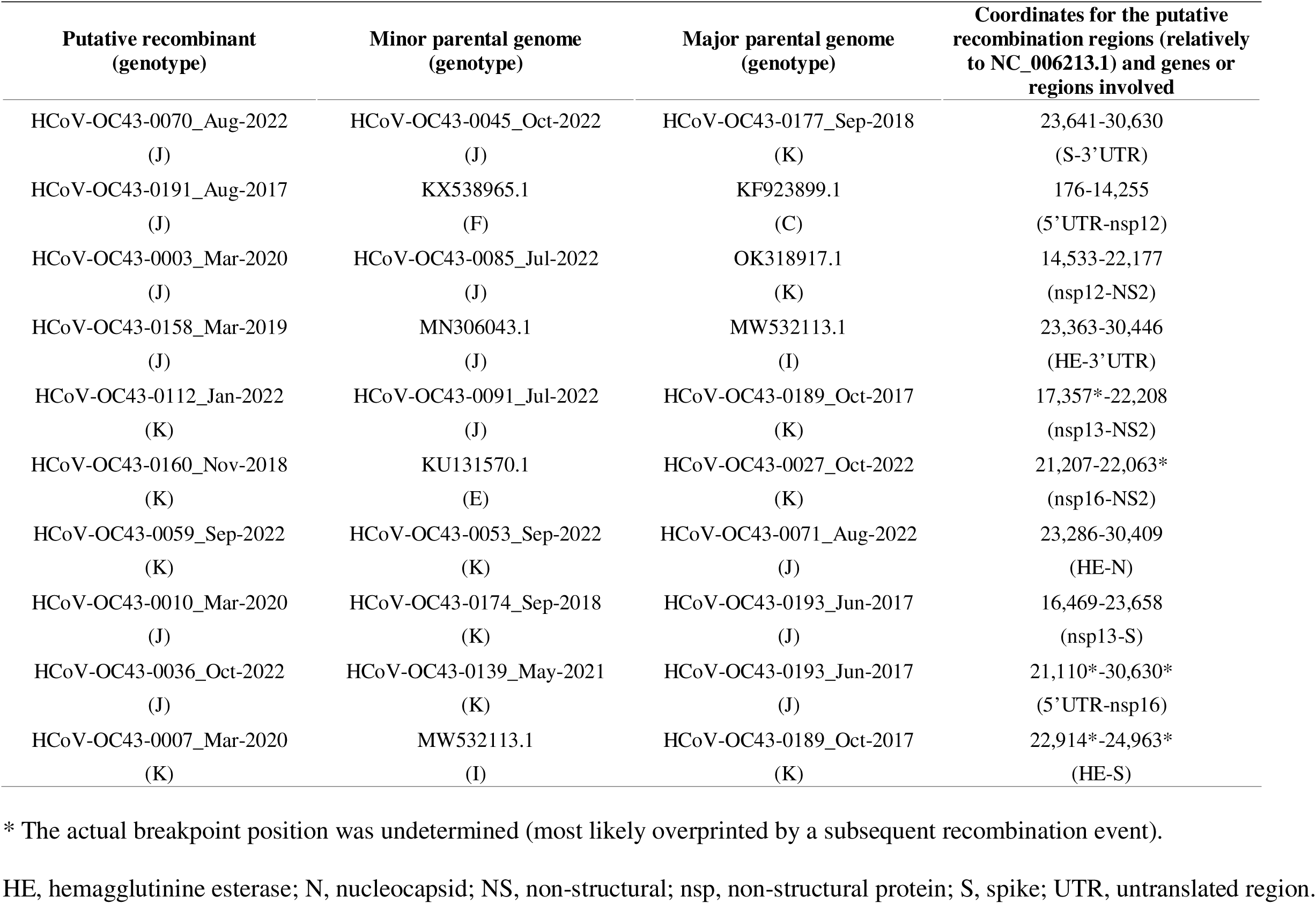
Putative HCoV-OC43 recombinant genomes detected and parental genomes either obtained here or collected from GenBank.

| Putative recombinant<br>(genotype) | Minor parental genome<br>(genotype) | Major parental genome<br>(genotype) | Coordinates for the putative<br>recombination regions (relatively<br>to NC_006213.1) and genes or<br>regions involved |
| --- | --- | --- | --- |
| HCoV-OC43-0070_Aug-2022<br>(J) | HCoV-OC43-0045_Oct-2022<br>(J) | HCoV-OC43-0177_Sep-2018<br>(K) | 23,641-30,630<br>(S-3'UTR) |
| HCoV-OC43-0191_Aug-2017<br>(J) | KX538965.1<br>(F) | KF923899.1<br>(C) | 176-14,255<br>(5'UTR-nsp12) |
| HCoV-OC43-0003_Mar-2020<br>(J) | HCoV-OC43-0085_Jul-2022<br>(J) | OK318917.1<br>(K) | 14,533-22,177<br>(nsp12-NS2) |
| HCoV-OC43-0158_Mar-2019<br>(J) | MN306043.1<br>(J) | MW532113.1<br>(I) | 23,363-30,446<br>(HE-3'UTR) |
| HCoV-OC43-0112_Jan-2022<br>(K) | HCoV-OC43-0091_Jul-2022<br>(J) | HCoV-OC43-0189_Oct-2017<br>(K) | 17,357*-22,208<br>(nsp13-NS2) |
| HCoV-OC43-0160_Nov-2018<br>(K) | KU131570.1<br>(E) | HCoV-OC43-0027_Oct-2022<br>(K) | 21,207-22,063*<br>(nsp16-NS2) |
| HCoV-OC43-0059_Sep-2022<br>(K) | HCoV-OC43-0053_Sep-2022<br>(K) | HCoV-OC43-0071_Aug-2022<br>(J) | 23,286-30,409<br>(HE-N) |
| HCoV-OC43-0010_Mar-2020<br>(J) | HCoV-OC43-0174_Sep-2018<br>(K) | HCoV-OC43-0193_Jun-2017<br>(J) | 16,469-23,658<br>(nsp13-S) |
| HCoV-OC43-0036_Oct-2022<br>(J) | HCoV-OC43-0139_May-2021<br>(K) | HCoV-OC43-0193_Jun-2017<br>(J) | 21,110*-30,630*<br>(5'UTR-nsp16) |
| HCoV-OC43-0007_Mar-2020<br>(K) | MW532113.1<br>(I) | HCoV-OC43-0189_Oct-2017<br>(K) | 22,914*-24,963*<br>(HE-S) |

\* The actual breakpoint position was undetermined (most likely overprinted by a subsequent recombination event).
HE, hemagglutinine esterase; N, nucleocapsid; NS, non-structural; nsp, non-structural protein; S, spike; UTR, untranslated region.

## DISCUSSION

As the SARS-CoV-2 pandemic that emerged in 2019 reminded us, coronaviruses are viruses that have a continuous potential of emergence both in non-human animals and in humans and are zoonotic source of infections for humans. Evidence of spill-over from humans to chimpanzees of HCoV-OC43 was also reported in Ivory Coast (Dinwiddle et al, 2016; Patrono et al, 2018). Therefore, it is critical to monitor the evolution of these viruses to account for the different mutants and variants observed. Among the four so-called ‘endemic’ coronaviruses that infect humans, HCoV-OC43 is the one the most frequently detected. Here, we obtained and analyzed 185 HCoV-OC43 genomes, whereas only 361 complete or near complete genomes were available in GenBank at the beginning of the present study (as of 04/10/2023). In addition, these GenBank sequences were obtained primarily in China and the USA, and only 11 were obtained in France. Moreover, for the 2017-2022 period studied here (as could be assessed when the respiratory sample collection date was available), there were a total of only 145 genomes available worldwide, and none were from France. Thus, here we expanded by approximately 50% the set of HCoV-OC43 genomes available worldwide until late 2023, we more than doubled the set of genomes available worldwide for the 2017-2022 period, and we expanded by approximately 17 times the number of genomes available for France and provided the only genomes for the period from 2017 to 2022 and this country.

PCR amplification of viral RNA pre-NGS is often necessary considering viral loads in clinical samples is often too low to obtain genomes, and even reads by direct NGS. This has been the case with SARS-CoV-2 during the pandemics with the implementation and updating of Artic systems (https://artic.network/ncov-2019/ncov2019-bioinformatics-sop.html). A protocol for whole-genome sequencing of HCoV-OC43 was previously described (Maurier et al, 2019). It was designed based on 47 previously available genomes and used 99 PCR primer pairs that generated 500-550 bp long amplicons that covered the whole genome but were pooled in 12 sets (each containing from 3 to 12 primer pairs). This protocol allowed obtaining 9 full-length or near full-length genomes from 11 respiratory samples. Another study conducted in Russia reported in 2025 the design using the PrimalScheme application (https://primalscheme.com/) of a panel of 36 PCR primers with a size of 900-1,100 bp for HCoV-OC43 whole genome amplification prior to NGS (Musaeva et al, 2025). Twenty-three genomes with >70% coverage were obtained, with the highest coverage observed in case of qPCR Ct of HCoV-OC43 RNA detection <25. Also, a study conducted in UK reported the design and use of an amplicon-based whole genome sequencing panel targeting all four endemic HCoV plus SARS-CoV-2 (McClure et al, 2025). PCR primers were approximately 1,200 bp-long. A total of 128 (69%) genomes with >95% coverage, which were of genotypes E, G, I, J and K, were obtained from 185 samples collected between 2016 and 2023. Here, PCR primer pairs were designed based on 220 complete genomes collected from GenBank, and 34 pairs were eventually retained that generated amplicons with sizes varying from 500 and 2,600 base pairs and were mixed into only two pools (of 17 primer pairs each). This protocol allowed obtaining 185 full-length or near full-length genomes from 434 nasopharyngeal samples with various Ct values. This PCR yield was only 43% but it must be taken into account that shorter genomes, which displayed a coverage <80% of the full-length reference genomes, were obtained from samples from which our protocol failed to generate larger genomes. It is worthy to note that another strategy to obtain whole HCoV-OC43 genomes in case of low viral load is an hybridization-based enrichment with an appropriate viral panel probe set, as was for instance reported in 2018 to obtain four genomes from respiratory samples of children with severe acute respiratory infections or to investigate using genomics an outbreak in wild chimpanzees in Ivory Coast (Dinwiddle et al, 2016; Patrono et al, 2018).

The predominance observed here of genomes obtained for the 2020-2022 period is due to the Covid-19 pandemic that was associated with an increase in the sampling of patients with clinical symptoms of respiratory infection. We identified three HCoV-OC43 genotypes, G, J and K. Genotypes J and K, which are among the latest detected HCoV-OC43 genotypes, were reported for the first time in China in 2022 (Zhang et al, 2022). Ye et al recently reported the detection of four genotypes (G, I, J and K) and of an emerging lineage (branching with genotypes I and K in the phylogenetic analysis) by analyzing 179 complete or near complete genomes obtained from 312 HCoV-OC43 RNA-positive respiratory samples collected in China (Ye et al, 2023). We show here that three of these four genotypes also circulated in France during the period of our study, and this finding questions on since how long they were circulating in our country. According to the period of time of our study, either a unique genotype was found to circulate (from autumn 2020 until autumn 2021) or alternatively two or three genotypes co-circulated. Otherwise, none of the genomes obtained here belonged to the new HCoV-OC43 lineage that was reported to had emerged in China (Ye et al, 2023).

The HE gene was the structural gene showing the greatest diversity for genotypes J and K genomes, while the envelope gene showed the greatest diversity for genotype G. The Nsp13 gene that encodes an helicase was the informational gene that exhibited the greatest diversity. The spike gene was one of those with the greatest diversity in the present study, which is in accordance with other studies showing that this gene evolves rapidly in coronavirus genomes (Lau et al, 2011; Ye et al, 2023). For instance, a considerable genetic diversity was previously reported based on clonal sequences of the S1 gene of seven clinical HCoV-OC43 (Vabret et al, 2006). Besides, Oong et al reported that spike mutations D267K and I268D (which correspond to D266K and I267D, respectively, in the study by Ren et al), were signature mutations of the genotype G (Ren et al, 2015; Oong et al, 2017). As a matter of fact, these mutations were not carried by genotype G genomes obtained here but only by some of the genotype G genomes from GenBank. Oong et al also reported the presence of mutations P22T and T271S in two genomes from China (Ren et al, 2015). All the genomes obtained here carried the mutation P22T, but some genomes available in GenBank did not. As for mutation T271S, it was harbored by some but not all genotype G genomes obtained here or from GenBank. Thus, the classification of HCoV-OC43 genomes based on signature mutations needs to be checked and refined based on currently available genomes as it depends on the number and diversity of available genomes. Finally, heere, one mutation located in the spike RBD, which was associated with capabilities to escape immune responses and adapt to humans (Hulswit et al, 2019), was identified in each of the three HCoV-OC43 genotypes G, J and K. Taken together, previous findings hence suggest that tracking HCoV-OC43 evolution based on genome analysis is worthy to gain a better picture and understanding of the emergence and faith of viral lineages within and between countries.

HCoV-OC43 is a coronavirus known to frequently recombine in both humans and other animals (Lau et al, 2011; Zhang et al, 2011; Zhu et al, 2018; Wells et al, 2023). Recombination between two different ‘parental’ genomes occurs when there is a coinfection of same host cell, resulting in the emergence of a virus that harbors parts of the genetic material of both ‘parents’. Several of the 11 currently defined HCoV-OC43 genotypes (A-K) were reported to be derived from recombination events or from a recombinant ‘parent’. Regarding genotype K, it was previously reported to result from a recombination involving a parental genome of genotype I ()Zhang et al, 2022), while genotype J was previously reported as being involved in recombination involving parental genomes of genotypes H and I (Zhang et al, 2022), and genotype H was suspected to have occurred from recombination between parental genomes of genotypes F/D-like, E and B (Zhu et al, 2018). In the present study, seven genotypes were putatively involved in 10 different combinations and putative recombinations involved six different genes and the two UTR regions at the tips of the HCoV-OC43 genome. Particularly, putative recombinants were predicted that involved genotypes K and J, which were those very majoritary in our dataset and which co-circulated during several periods of time. Seven of the ten putative recombinant genomes were comprised by a genotype J fragment as major or minor parental genome in three and four cases, respectively.

In summary, the present study demonstrates the circulation in France between 2017 and 2022 of new HCoV-OC43 genotypes initially detected in China, and deciphered their dynamics over time. It contributes to start filling a huge gap on HCoV-OC43 genome availability worldwide and in France, and points out the tremendous imbalance between the amounts of genomic data on SARS-CoV-2 and on another human coronavirus discovered three decades earlier and known to cause frequent and sometimes severe infections in humans. Novel HCoV-OC43 genotypes have been recurrently reported (Lau et al, 2011; Zhang et al, 2015; Oong et al, 2017; Zhu et al, 2018; Zhang et al, 2022) including some involved in severe and fatal cases (Zhang et al, 2022; Lau et al, 2022), and this suggests that new variants will likely emerge in the near future. Besides, previously unannotated open reading frames (ORFs) were recently reported in the HCoV-OC43 genome including a putative nested ORF as well as short upstream ORFs, which warrants investigating for various isolates belonging to circulating genotypes the viral transcriptional and translational landscapes (Bresson et al, 2025). Finally, further studies are needed to gain a better knowlege of whether the different HCoV-OC43 genotypes are associated with particular transmissibility, antigenicity, replicative capacity, or pathogenicity, and of the determinants of the rise and fall of their epidemics.

## Acknowledgments

We are thankful to the technical team of the IHU Méditerranée Infection next-generation sequencing platform.

## Data Availability

The HCoV-OC43 genomes analysed in this study were submitted to the Genbank database (https://www.ncbi.nlm.nih.gov/genbank/) (GenBank Accession no. PV471628-PV471812).

## Author contributions

Conceived and designed the experiments: BLS, PC. Contributed materials, analysis tools: All authors. Analyzed the data: HH, CB, LL, BLS, PC. Writing—original draft preparation: HH, BLS, and PC. Writing—review and editing: All authors. All authors have read and agreed to the published version of the manuscript.

## Ethics

The present study has been registered on the Health Data Access Portal of Marseille public and university hospitals (Assistance Publique-Hôpitaux de Marseille (AP-HM)) with No. PADS24-190 and was approved by the Ethics and Scientific Committee of AP-HM.

